# The Atomic Structure of the Microtubule-Nucleating γ-Tubulin Small Complex and its Implications for Regulation

**DOI:** 10.1101/310813

**Authors:** Axel F. Brilot, David A. Agard

## Abstract

The microtubule cytoskeleton is essential in mediating a number of critical cellular processes, affecting cell shape, transport, organelle organization, and chromosomal segregation during mitosis. Microtubule network dynamics are controlled by many factors including the efficiency and localization of the nucleation machinery. Microtubule nucleation is dependent on the universally conserved γ-tubulin small complex (γTuSC), a 300 kDa heterotetramer composed of two copies of γ-tubulin and one each of accessory proteins GCP2 and GCP3. In yeast, nucleation is mediated by a heptameric ring of γTuSC, which presents 13 γ-tubulins to form a template for microtubule nucleation.

We have obtained single-particle structures of the γTuSC as a monomer and dimer at resolutions of 3.6-4.6Å, allowing us to build an atomic model for this important complex. By comparison with a crystal structure of isolated γ-tubulin, it is clear that γ-tubulin is activated upon assembly into the γTuSC, in a manner analogous to the bent to straight transition in αβ-tubulin upon assembly into the microtubule lattice. Our structures allow us to map phosphorylation sites, revealing several at key interfaces, highly suggestive of their role in regulating spindle pole body attachment and assembly into rings. When combined with previous lower resolution structures of helical assemblies, we observe that additional conformational changes occur during ring assembly and activation.

## Introduction

The microtubule (MT) cytoskeleton plays an essential role in the spatio-temporal control of eukaryotic cellular organization, cytoplasmic transport and chromosome segregation during mitosis(Desai and Mitchison, 1997). The organization and function of the cytoskeletal network is tightly controlled by regulating the rate and location of nucleation, as well as MT polymerization kinetics and stability(Akhmanova and Steinmetz, 2015; Howard and Hyman, 2009; Teixidó-Travesa et al., 2012).

In most cells, MT nucleation occurs primarily at MTOCs and is dependent on the universally conserved γ-tubulin ring complex (γTuRC)(Luders and Stearns, 2007). In metazoans and plants, the γTuRC is recruited to MT nucleation sites as a large, pre-formed ring-shaped 2.2 MDa complex (Teixidó-Travesa et al., 2012). Although the exact stoichiometry of the metazoan γTuRC is unknown, it is composed of many copies of γ-tubulin, and a smaller number of the γ-tubulin binding proteins, GCP2-6, as well as other accessory proteins. GCP2 and GCP3 form a stable 300 KDa subcomplex with 2 copies of γ-tubulin (the γTuSC), and are present in multiple copies(Murphy et al., 2001; Oegema et al., 1999). While only ~15% homologous to one another and varying in size from 70KDa to 210 KDa, GCP2-6 share a conserved core of 2 grip domains(Kollman et al., 2011). Structural and biochemical studies have shown that the N-terminal grip1 domain drives lateral association between GCPs, while the C-terminal grip2 domain binds to γ-tubulin(Choy et al., 2009; Farache et al., 2016; Greenberg et al., 2016; Guillet et al., 2011; Kollman et al., 2015).

In budding yeast, only homologues of GCP2and GCP3 (Spc97p and Spc98p) and γ-tubulin (Tub4p) are present (Vinh et al., 2002). Spc110p, a distant pericentrin homologue, recruits this complex to the nuclear face of the spindle pole body (SPB), while Spc72p recruits it to the cytoplasmic face(Knop and Schiebel, 1997, 1998; Nguyen et al., 1998).

Previous moderate resolution cryoEM structural studies (8Å) had shown that γTuSCs complexed with the N-terminal domain of Spc110p self assemble into filaments having 6.5 γTuSCs/turn thereby presenting 13 γ-tubulins to template 13-protofilament MTs (Kollman et al., 2010, 2015). Although close to MT symmetry, the γ-tubulins within each γTuSC were too far apart to correctly match the MT lattice. The relevant *in vivo* conformation was determined by cryo-tomography and sub-volume averaging, clearly showing a MT-matching geometry at the yeast spindle pole body, suggesting that γTuSC closure might be an important regulatory step (Kollman et al., 2015). To validate this hypothesis, γ-tubulin was engineered with disulfides to stabilize a closed MT-like conformation, resulting in significantly enhanced MT nucleation (Kollman et al., 2015). This also had the benefit of improving the cryoEM map (6.5Å) to a point where it became feasible to build an initial pseudoatomic model (Greenberg et al., 2016; Kollman et al., 2015) based on the crystal structure of human γ-tubulin (Aldaz et al., 2005; Rice et al., 2008) and the distant and much smaller human Gcp4 (Guillet et al., 2011; Kollman et al., 2015).

Biochemical studies on the role of Spc110p, revealed that higher order oligomerization of Spc110p was required for γTuRC assembly at physiologically relevant concentrations of γTuSC (Kollman et al., 2010, 2015; Lyon et al., 2016). Together with the structural data, this suggested that regulation of both γTuSC conformation and γTuRC assembly would be important for ensuring that proper MT formation only happened at the spindle pole body. Supporting the need for precise regulation of γTuRC assembly and function, all the components of the γTuSC, as well as Spc110p and Spc72p are phosphorylated in a cell-cycle dependent manner(Keck et al., 2011). Many of these phosphorylation events, particularly in γ-tubulin or Spc110p, have been implicated in cellular viability, spindle morphology, or shown to have effects on γTuSC assembly (Huisman et al., 2007; Keck et al., 2011; Lin et al., 2011, 2014; Lyon et al., 2016; Peng et al., 2015; Vogel et al., 2001). However, the mechanism of γTuRC assembly and activation and its regulation by phosphorylation or other factors remains unclear.

Using single particle cryoEM, we present here maps of monomeric and dimeric γTuSCs at near atomic resolution. These have allowed de-novo model building of unknown regions and reinterpretation of significant regions of γTuSC structure. Many of the annotated phosphorylation sites had fallen in regions of γTuSC not represented in the previous homology model, thus the new structure provides a powerful atomic framework for understanding the importance and mechanism of regulatory modifications. The high-resolution structure also reveals conformational changes that occur within γ-tubulin when it assembles into a γTuSC and some key origins for the remarkable 100-fold species preference of yeast γTuRCs for yeast tubulin.

## Results

### Cryo-Electron microscopy structure of the γTuSC

To resolve fundamental questions on the molecular basis for MT nucleation by γTuRC, as well as its assembly and activation, we determined the cryo-EM structure of γ-TuSC from images of frozen-hydrated single particles. At the concentration of ~1 μM used in data collection, micrographs and 2D classes show a mixture of γ-TuSC monomers and dimers, with a small number of larger oligomers (Fig. S1). We were able to obtain structures of the γ-TuSC monomer at ~3.6-4.1Å, and of a γ-TuSC dimer at ~4.6Å resolution using a ~1.3 million particle dataset (Figure 1,S2). An atomic model was built into these maps using Rosetta (DiMaio et al., 2015; Frenz et al., 2017; Wang et al., 2016) by combining information from the previously published pseudoatomic model of the γ-TuSC(Greenberg et al., 2016), the x-ray structure of human GCP4(Guillet et al., 2011), and the x-ray structure of γ-tubulin(Rice et al., 2008). Missing and incorrect regions were either manually corrected or sampled using Rosetta prior to further refinement. We were able to fit 2197 of 2615 residues present in the structure in our most complete models, accounting for ~84% of the molecular weight of the γTuSC. In these models, the C-terminal 20 and 27 residues of the γ-tubulins bound to Spc98p and Spc97p respectively, the N-terminal 179 residues and a 17-residue insertion of Spc98p, as well as the N-terminal 51 amino-acids, 110 amino acids in 4 insertions and the C-terminal 14 residues of Spc97p are not modelled.

**Figure 1:**
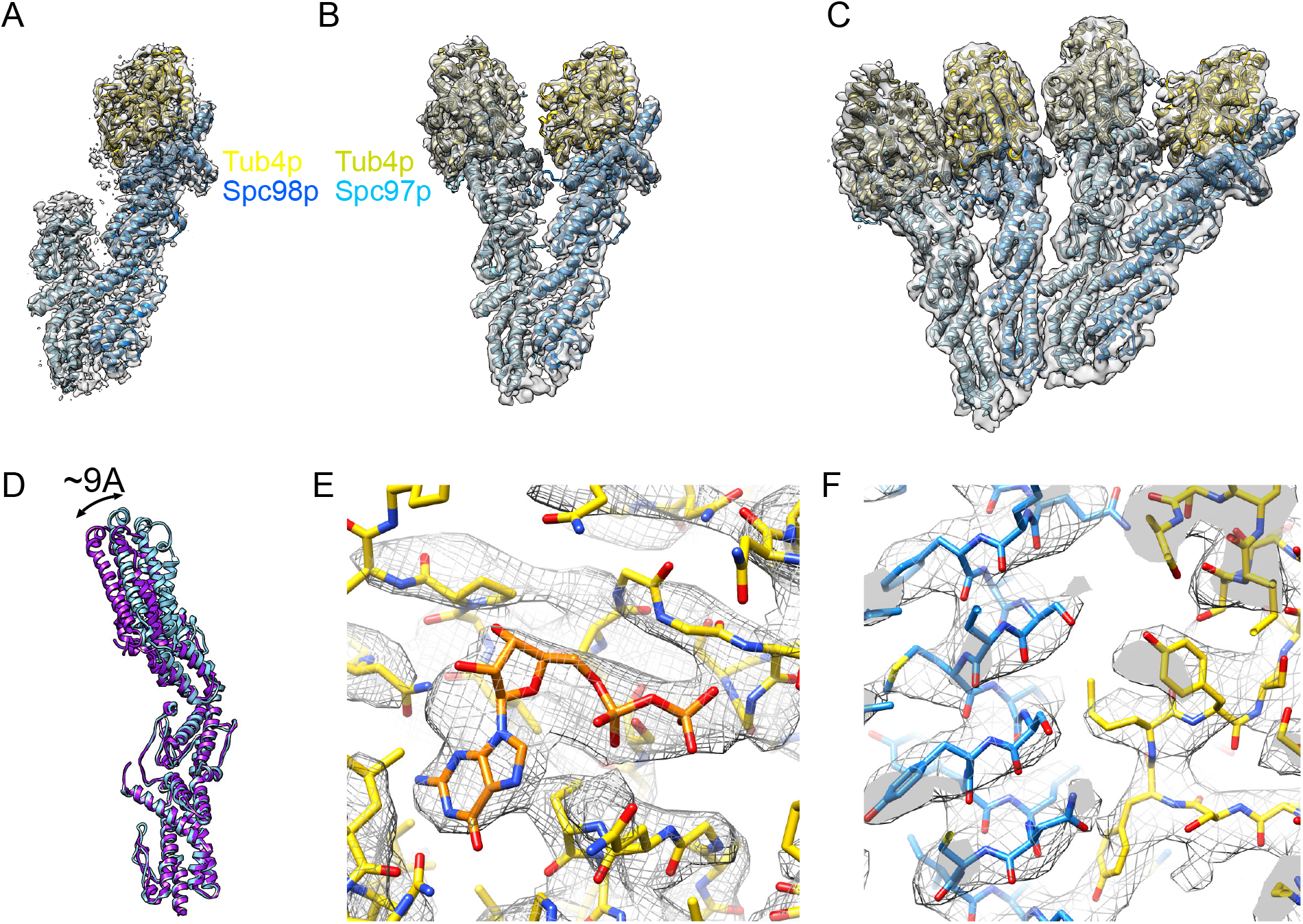
Single Particle Reconstructions of the γTuSC. A. ~3.6 Å reconstruction obtained by masking the mobile C-terminus of Spc97p showing the N-terminus of Spc97p (sky blue), Spc98p (dodger blue) and *γ*-tubulin bound to Spc98p (gold) B. ~3.9 Å reconstruction of gamma-TuSC colored as in panel A, with gamma-tubulin bound to Spc97p colored in khaki. C. ~4.6 Å reconstruction of a dimer of γTuSC colored as in panel B. D. View of Spc97p showing a ~9 Å motion of the C-terminus when the structure are aligned using Spc98p and its associated gamma-tubulin. E. Unmasked density of the GDP binding site of *γ*-tubulin in the ~3.6 Å structure, with GDP shown in orange. F. Unmasked density of the interface between *γ*-tubulin and Spc98p in the ~3.6 Å structure, colored as in Figure 1A.

The monomer structure has a characteristic V-shape, ~170Å in height and ~120Å wide. The N-terminal grip domains of GCP2 and GCP3 form the base of the structure, while the C-terminal grip domain contacts the γ-tubulins, which are positioned next to each other in an open conformation. The γ-TuSC dimer is formed of two γ-TuSCs in lateral contact using the same interface as observed in the γ-TuSC:Spc110p filament structures(Kollman et al., 2010, 2015).

3D classification of γ-TuSC monomers shows an ~9Å correlated motion in solution of the C-terminal grip2 domain of Spc97p and its associated γ-tubulin relative to Spc98p and its associated γ-tubulin in solution. This motion is consistent with previous work showing γ-TuSC is a highly dynamic molecule, with the GCPs capable of conformational changes resulting in large motions of the γ-tubulins (Greenberg et al., 2016; Kollman et al., 2008, 2010, 2015). Thus, our highest resolution reconstruction was obtained by filtering this region of density in the reconstructions to a lower resolution during alignment and reconstruction as implemented in Frealign (Grigorieff, 2016) (Figure 1A, S2).

### γ-Tubulin is activated upon assembly into the γ-TuSC

It has long been appreciated that αβ-tubulin heterodimers undergo a bent to straight transition upon microtubule assembly. The bent state is traditionally observed in complexes with GDP, drugs or proteins that inhibit microtubule polymerization, while the straight state has only been observed in microtubules(Alushin et al., 2014; Nawrotek et al., 2011; Ravelli et al., 2004). The bent to straight state conformational change in αβ-tubulin is characterized by a motion of H6, the H6-H7 loop and H7 towards the (-)-end, when the tubulins are aligned using the N-terminal domains (Figure 2A). Upon straightening, the H6-H7 loop moves several Angstroms, remodeling and strengthening the longitudinal interface, and relieving steric clashes that would have occurred with the C-terminal domain of the subsequent α-tubulin in a microtubule. Surprisingly, in solution GTP fails to drive a bent-straight conformational change, indicating that it is microtubule lattice and not the GTP that serve as the allosteric effector (Rice et al., 2008).

**Figure 2:**
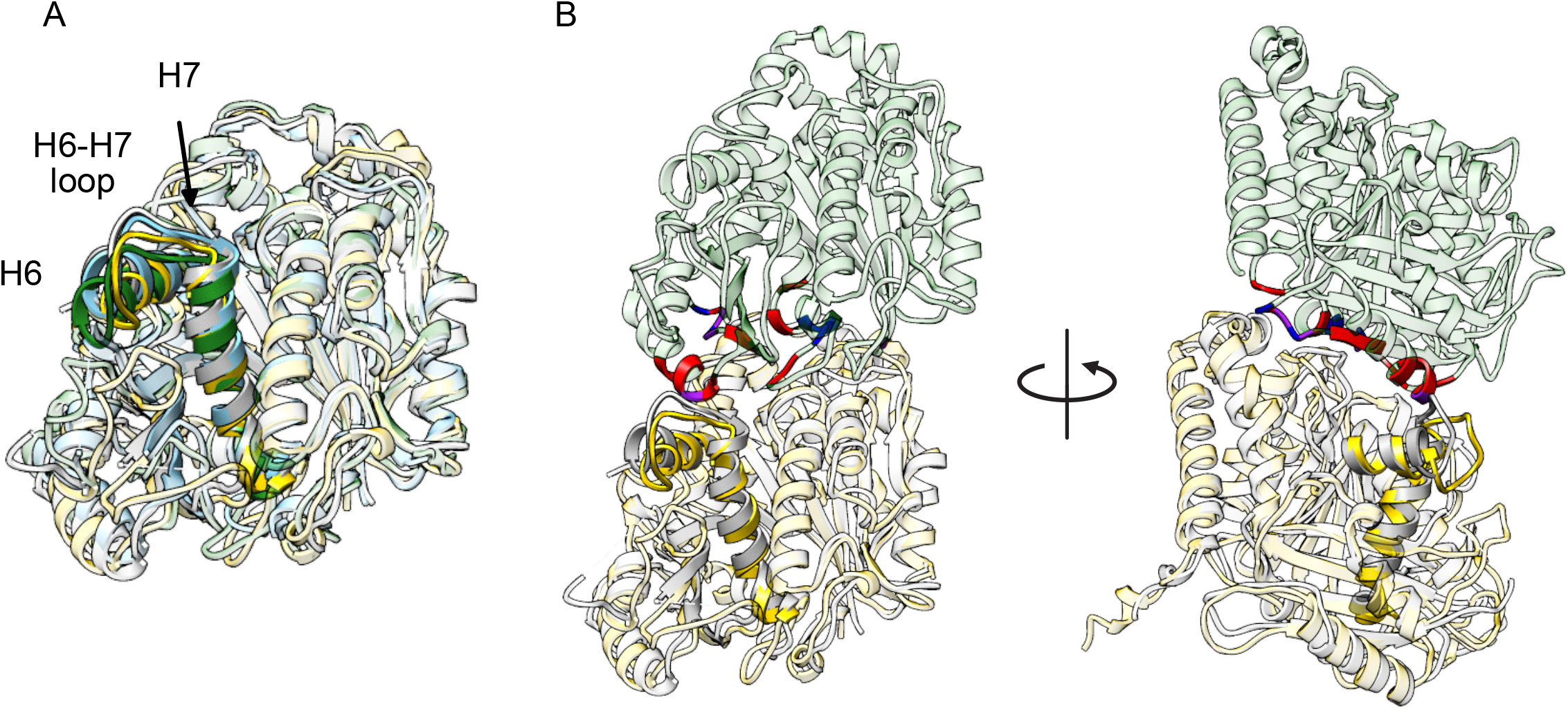
γ-Tubulin is Activated Upon Assembly Into The γ-TuSC. A. An overlay of yeast straight β-tubulin (forest green, PDB ID 5W3F) γ-tubulin bound to Spc98p (gold, this work), yeast bent β-tubulin (sky blue, PDB ID 4FFB) and the human free γ-tubulin (gray, PDB ID 3CB2) structures shows that the γ-tubulin bound to Spc98p H6-H7 loop adopts a conformation similar to that observed in straight microtubules. B. Clashes between γ-tubulin bound to Spc98p (blue) or human γ-tubulin (red, PDB ID 3CB2) or both (purple) with a straight α-tubulin are illustrated on a forest green α-tubulin. The yeast microtubule structure (PDB ID 5W3F) was used to model the βα-tubulin interface, and other proteins were aligned by the N-terminal domains. Clashes with a greater than 1.2 Å overlap are shown.

Previous high-resolution structures of γ-tubulin have shown it adopts a bent state when not in complex with GCPs independent of its nucleotide state. This raised the question if it too could change upon assembly into γTuSC. In our structures, γ-tubulin adopts a conformation distinct from the previously observed bent or straight conformations of yeast tubulin (Figure 2A. While H6, the H6-H7 loop and H7 are displaced relative to the positions observed in bent β-tubulin, the C-terminus of H6 is displaced laterally towards the N-terminal domain, relative to the axis observed in the bent to straight transition of yeast β-tubulin. However, despite these differences, the H6-H7 loop adopts a conformation strikingly similar to that observed in straight β-tubulin (Figure 2A), indicating that conformational changes of the γ-tubulin do occur upon assembly into γTuSC and likely serve to activate its function in microtubule nucleation. To examine this in more detail, we aligned the yeast γ-tubulin from our structure with a β-tubulin within the yeast MT (Howes et al., 2017) to simulate a γ-tubulin: α-tubulin (γTuSC:microtubule) interface (Fig 2B). This revealed that the previously observed bent γ-tubulin conformation would cause a significant number of steric clashes, while the majority of these were alleviated when *γ*-tubulin is assembled into the γTuSC (Figure 2B). Interestingly, the T5 loop, the other region in the γ-tubulin crystal structure that would clash, is a region of significant sequence difference between human and yeast γ-tubulins, suggesting an explanation for the very strong species preference observed experimentally (Kollman, 2015).

### γ-tubulin-γ-GCP interface

*γ*-tubulin forms an extended interface with the C-terminal grip2 domain of Spc97p and Spc98p (Figure 3). Contacts with the *γ*-tubulin N-terminal domain are primarily with the small domain adjacent to the two C-terminal helical bundles, with the small domain of Spc98p contacting the base of S1 and the H1-S2 loop. The C-terminal helical bundles contact the C-terminal domain of *γ*-tubulin forming extensive contacts with helix 8, the H8-S7 loop, s9, and helix 10. The H7-H8 loop is pinned between the H12-H13 loop at the N-terminus of the small domain and the C-terminal helical bundles. Helix 21, which in the GCP4 structure was docked against helices 19 and 20, adopts a different conformation between helix 20 of the GCPs, and helix 10 of *γ*-tubulin. The C-terminal residues of the *γ*-tubulin attached to Spc98p adopt a helical conformation, contacting helices 16 and 20 of Spc98p. Electrostatic interactions or salt bridges appear to play a significant role in defining the interface between the GCPs and *γ*-tubulin, as a significant portion of the interface region is composed of acidic residues on *γ*-tubulin and basic residues on the GCPs (Figure 3CD).

**Figure 3:**
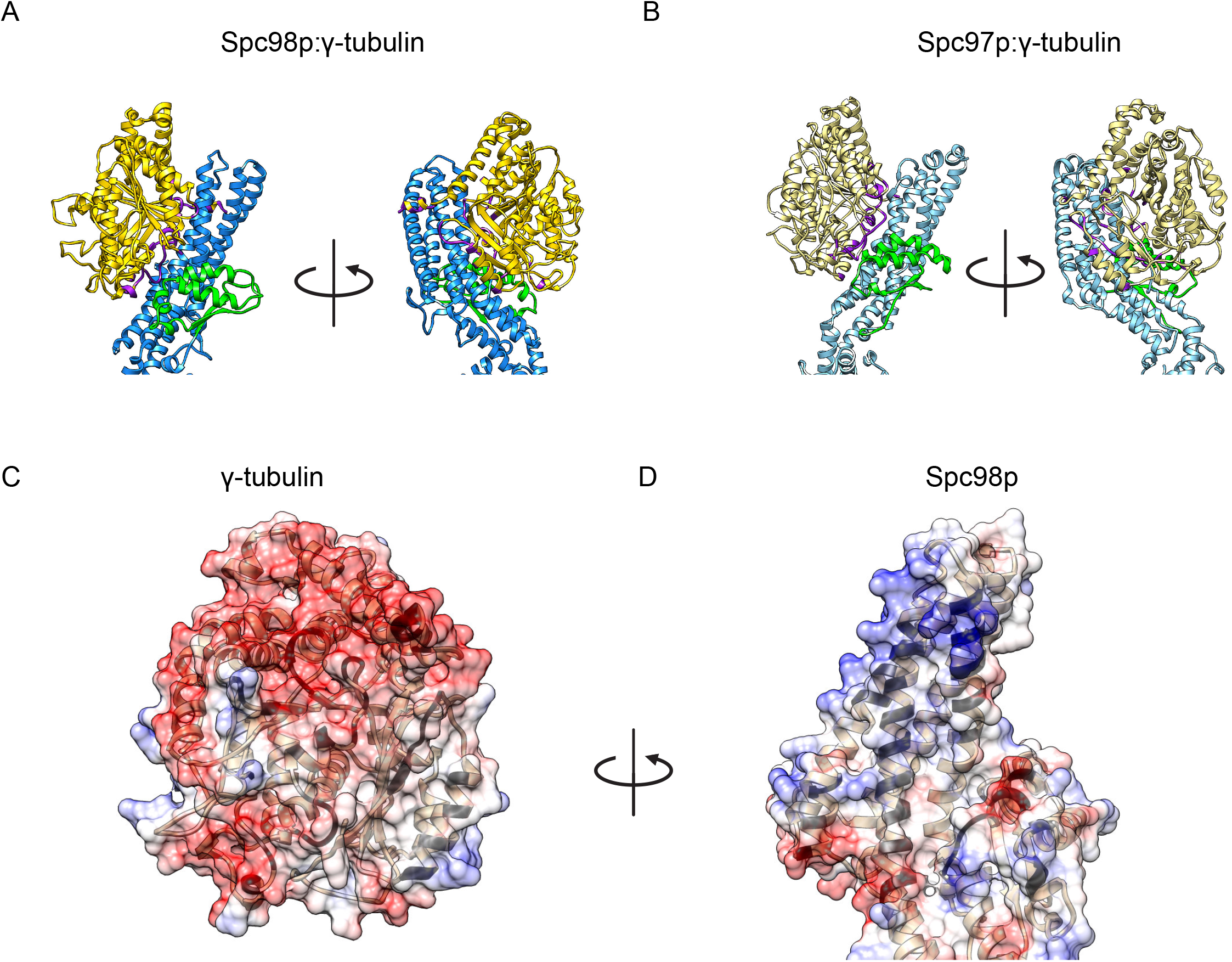
The γ-tubulin GCP binding interfaces. A. Close up views of the Spc98p γ-tubulin interface highlighting contacts between two proteins,. Spc98p is colored dodger blue, except for the small domain adjacent to the C-terminal helical bundles colored green. γ-Tubulin is colored in gold, with residues at the contact interface colored purple. B. Close up views of the Spc98p γ-tubulin interface highlighting contacts between two proteins,. Spc97p is colored light blue, except for the small domain adjacent to the C-terminal helical bundles colored green. γ-Tubulin is colored in khaki, with residues at the contact interface colored purple. C. A view of the Spc98p binding surface of *γ*-tubulin, colored according to Coulombic potential (calculated in UCSF Chimera). Contact residues are highlighted in black on tan. D. A view of the γ-tubulin binding surface of Spc98p, colored according to Coulombic potential (calculated in UCSF Chimera). Contact residues are highlighted in black on tan.

### Intra- and inter-γTuSC interfaces

The monomer interface between the GCPs is largely formed from the 2 N-terminal helical bundles of Spc97p and Spc98p (Figure 4A). The N-terminal interface has a prominent hydrophobic core involving residues from helix H1, the loop N-terminal to helix 1, the N-terminus of helix H2 and helices H3 and H4 on Spc97p (numbered by structural homology to GCP4 (Guillet et al., 2011)), with additional contacts being formed between the H7-H8 loop and the N-terminus of H2, the H2-H3 loop, and the C-terminus of H3, as well as the loop N-terminal to helix 8.

**Figure 4:**
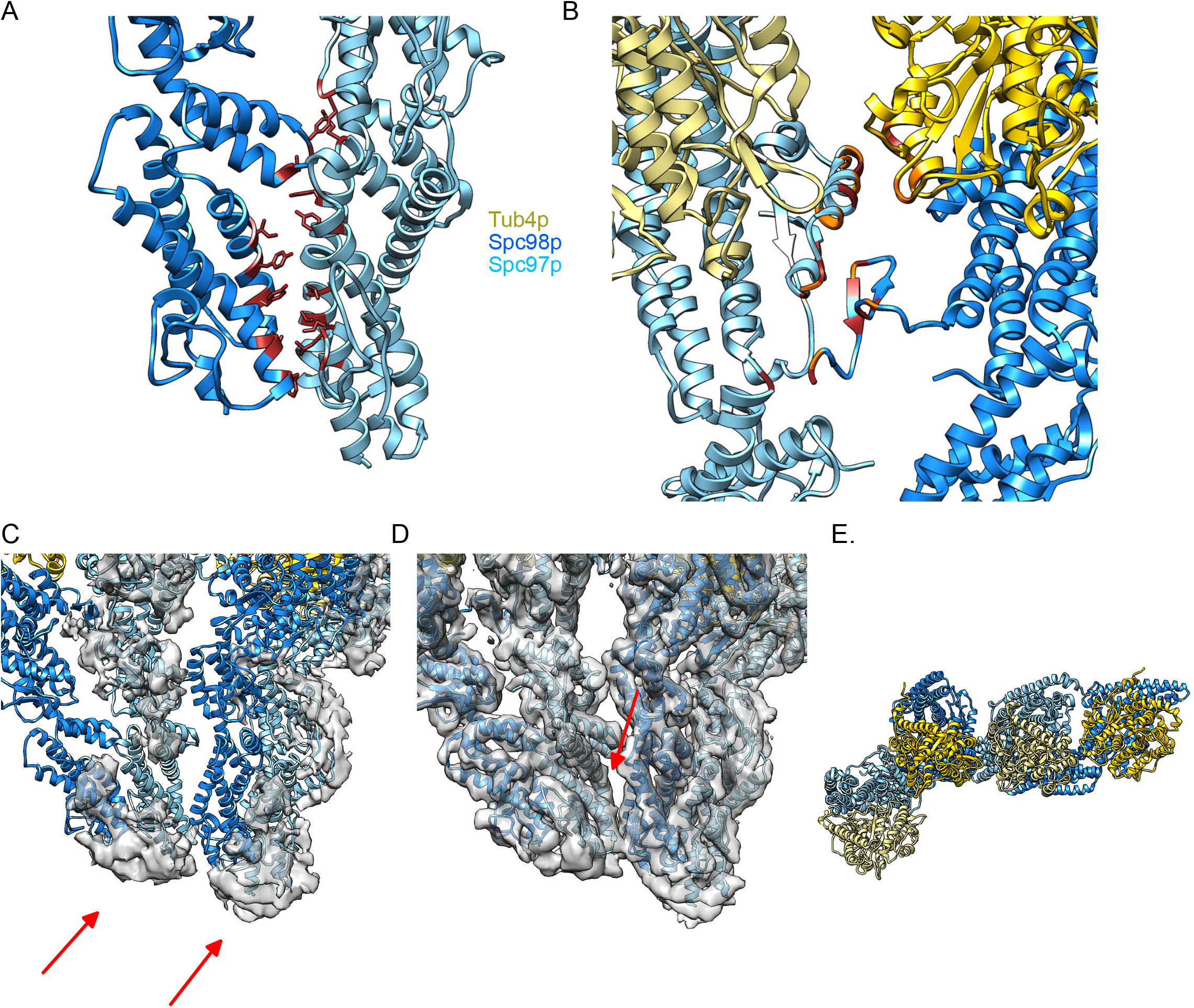
The γTuSC Monomer and Dimer Interfaces. A. Close up view of the N-terminal γTuSC monomer interface, colored as in Figure 1B, showing hydrophobic residues at the interface in brown. B. Close up view of the N-terminal γTuSC monomer interface, colored as in Figure 1B, showing hydrophobic residues at the interface in brown, and hydrophilic or charged residues in orange. C. The N-terminus of the γTuSC dimer, colored as in Figure 1C, overlaid with a difference map between the map and model, showing significant contiguous density at the N-terminus (as indicated by the arrows) indicating unassigned density contributing to contacts with each monomer at the N-terminus. Noise in the difference map is hidden using the “Hide Dust” function in UCSF Chimera. D. ~4.6 Å reconstruction of the γTuSC dimer, colored as in Figure 1C, as seen from the exterior face of an assembled *γ*TuRC, overlaying the N-terminus of Spc97p and Spc98p highlighting the differences in the packing at the monomer and dimer interfaces. The arrow highlights the loose packing of the dimer interface. E. Top view of the γTuSC dimer, colored as in Figure 1C.

A previously unmodeled 33 residue insertion forms contacts with strands B5 and B6 and helix 14 as well continuing through the interface between the third and fourth helical bundles at the monomer interface between Spc97p and Spc98p (Figure 4B).

A hydrophilic interface is formed between Spc97p helix 14 and helices 9, 10, as well as the S9-S10 loop of the *γ*-tubulin bound to Spc98p (Figure 4B). Significant unmodeled density on the external face of the *γ*TuSC highlighted in a difference map (Figure 4C) likely corresponding to the N-termini of Spc97p or Spc98p appears to form additional contacts at the monomer interface between Spc97p and Spc98p.

By contrast, simple inspection of the dimer interface reveals a much more loosely packed interface (Figure 4D). The dimer interface is composed of the opposite surface of the three N-terminal helical bundles of Spc97p and Spc98p. There are two main contact zones: the H2-H3 loop of spc98p forms contacts with the H7-H8 loop of Spc97p; helix H1, the loop N-terminal to it, and the C-terminus of H4 on Spc98p form contacts with the N-terminal region of H2 and H5 on Spc97p. (Figure 4D). The main contacts of the *γ*-tubulins at the dimer interface are with each other (Figure 4E) and appear similar to those observed in the open *γ*-tubulin helical structure(Kollman et al., 2010). Proximal to the *γ*-tubulins, a contact is made between helix 14 of Spc97p (GCP4 numbering) and the loop between helices 20 and 21 (Figure 4E).

### Twisting of α-helical bundles in Spc97p and Spc98p accommodate conformational changes of γ-TuSC during assembly

To assess changes that occur when two γTuSC monomers associate to form the observed dimer, the solvent exposed inter-γTuSC Spc97p:98p and the central inter-γTuSC Spc97:98p were aligned with the monomer Spc97:98p using helices H2-6 (residues 214-299) for Spc98p and helices H2-5 (residues 100-180) for Spc97p, which form the core of their 2 N-terminal helical bundles, and form an axis which roughly parallels the long axis of GCP2-3. From this it can be seen that the outer GCP2,3 rotate asymmetrically (Figure 5AB). Within each GCP, the middle and C-terminal helical bundles appear to undergo a twisting motion relative to the axis defined by the 2 N-terminal helical bundles (Figure 5AB). Comparing the monomer and dimer Spc98p structures shows that the relative twist is significantly less in the outer Spc98p of the dimer for Spc98p subunits involved in the dimer interface.

**Figure 5:**
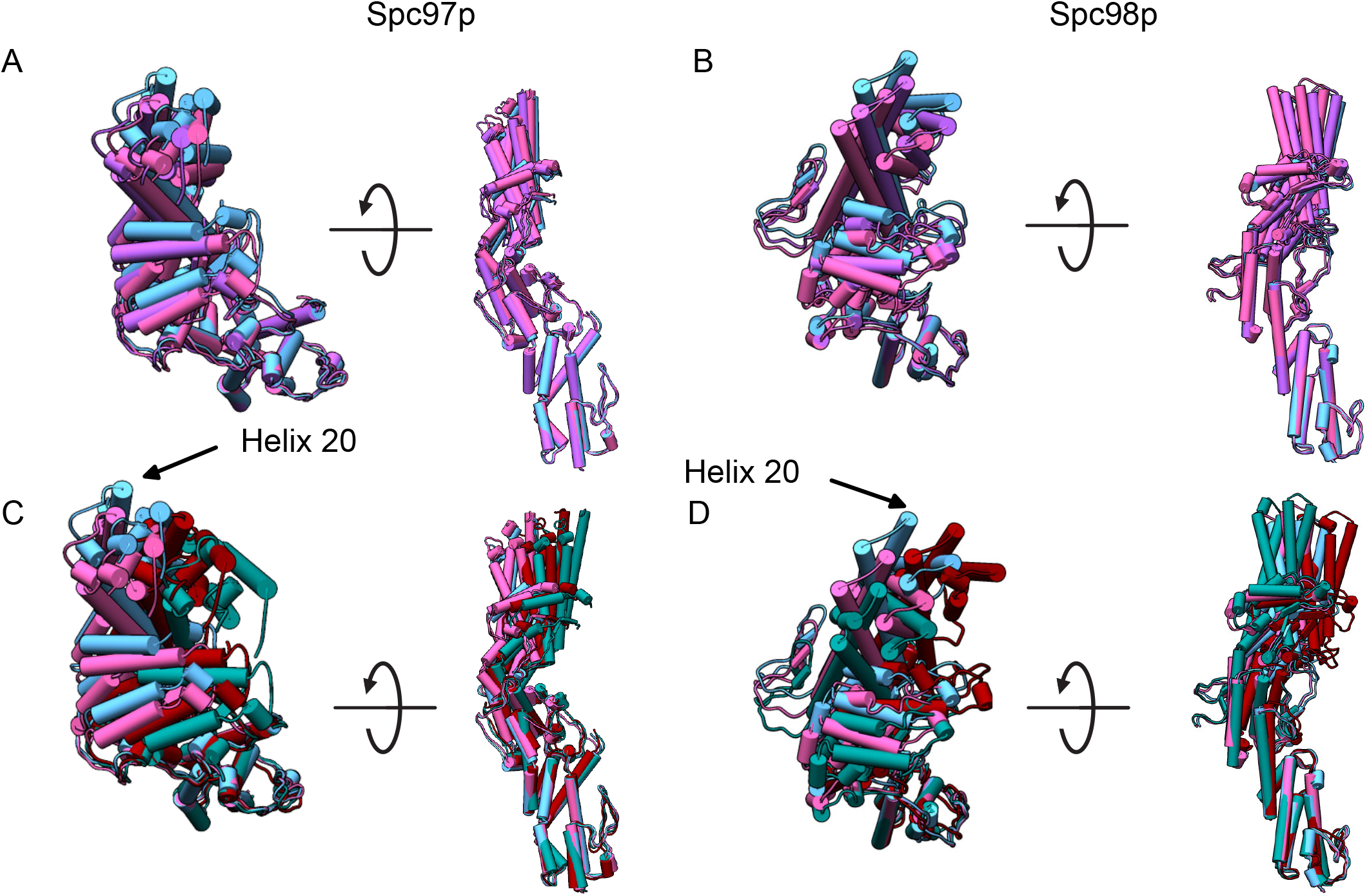
Conformational Changes of Spc97p and Spc98p During Assembly and Activation. A. Views of Spc97p aligned using the N-terminal helical bundles of the monomer (helices 2-5 GCP4 numbering, residues 100-180) (light blue), non-interface dimer (violet) and interface dimer (light pink) illustrating the twist of the helical bundles in our monomer and dimer models. B. Views of Spc98p aligned using the N-terminal helical bundles of the monomer (helices 2-6 GCP4 numbering, residues 214-299), non-interface dimer and interface dimer (colored as in A) illustrating the twist of the helical bundles in our monomer and dimer models. C. Views of Spc97p aligned using the N-terminal helical bundles of the monomer (light blue), interface dimer (light pink), open state helical structure (maroon) and closed state helical structure (bluish green) illustrating the twist of the helical bundles during assembly. D. Views of Spc98p aligned using the N-terminal helical bundles of the monomer, interface dimer, open state helical structure and closed state helical structure (colored as in C) illustrating the twist of the helical bundles during assembly.

Thus, even in the absence of Spc110p, lateral assembly of the γTuSCs requires remodeling of the GCPs.

To complement our structures of the monomer and dimer state of the γTuSC, we fitted our monomer state into the previously determined helical structures of the open and closed assembled states of the γTuSC in the presence of Spc110p(Kollman et al., 2010, 2015). While these structures are at lower resolution, precluding an atomic interpretation, these fits should inform on the overall conformation of the assembled open and closed states.

Comparing the monomer, interface, open state, and closed state Spc97p and Spc98p conformations shows these also undergo large twisting motions (Figure 5CD). The center of mass of helix 20 in Spc97p moves by ~20 Å, with the extremes occupied by the monomer conformation and the closed state conformation. Interestingly, the Spc98p at the dimer interface and that observed in the open state are twisted in opposite directions relative to the monomer Spc98p, indicating that either assembly or binding to Spc110p induces large conformational changes in Spc98p. The largest conformational change in Spc98p is observed between the open and closed states of Spc98p, with helix 20 moving ~20 Å.

### Mapping phosphorylation sites on the γTuSC suggests regulatory roles

The *γ*TuSC is phosphorylated in a cell-cycle dependent manner, and perturbations affecting its phosphorylation have been shown to affect spindle morphology (Fong et al., 2018; Keck et al., 2011; Lin et al., 2011; Peng et al., 2015; Vogel et al., 2001) To better understand the potential role of phosphorylation in γTuSC regulation and function, we mapped a recently determined set of phosphorylation sites, including a re-analysis of previously determined data and newly acquired data from SPBs(Fong et al., 2018), onto our monomeric *γTuSC* structure (Figure 6A).

**Figure 6:**
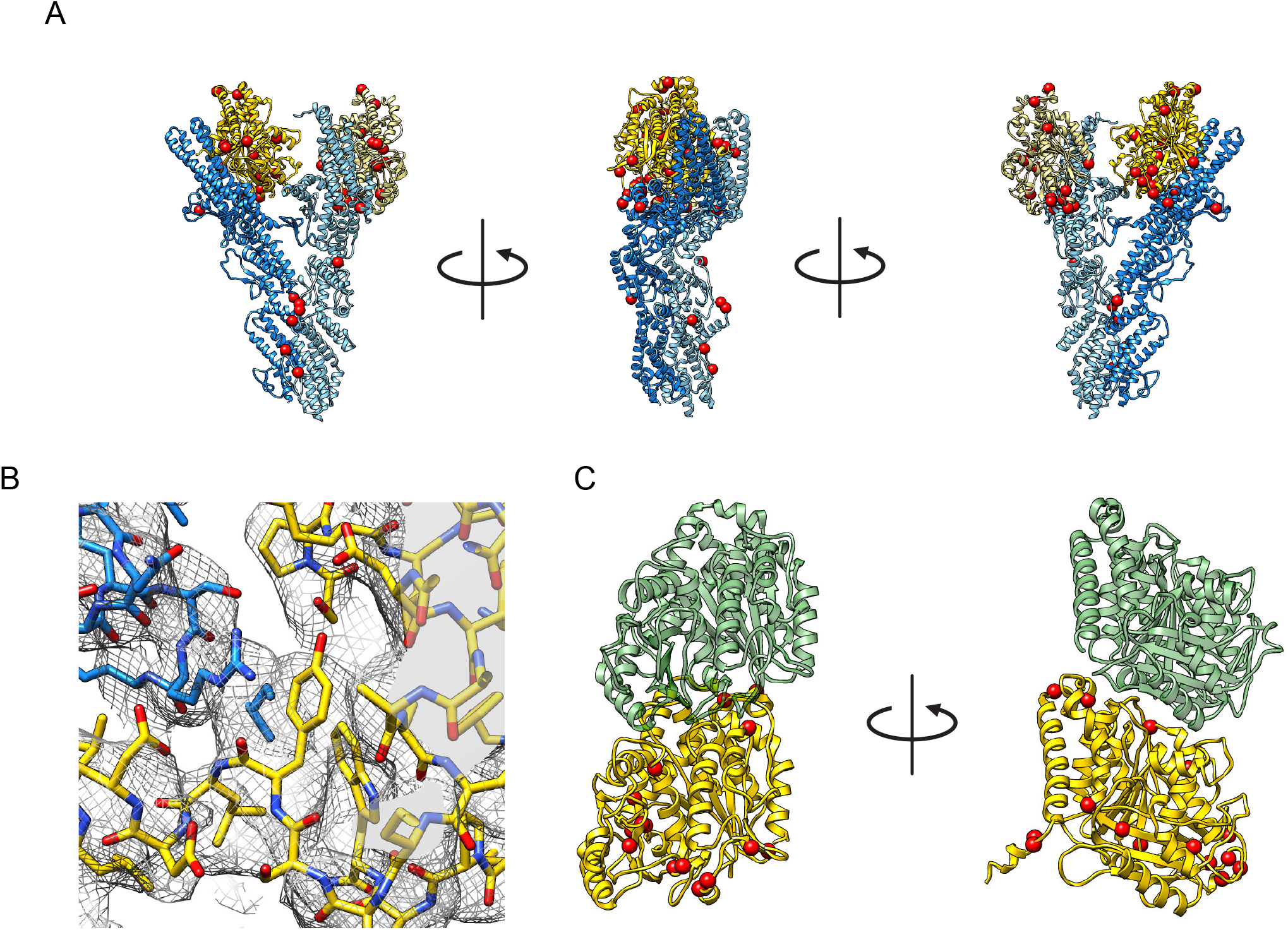
Phosphorylation sites visualized on the γTuSC structure. A. Three views rotated by 90 degrees of γTuSC with phosphorylation sites marked with red ball. Ribbons are colored as in Fig. 1B B. View of the contact between γ-tubulin Y445 and Spc98p R633 at the interface between the C-terminus of γ-tubulin and Spc98p. C. Phosphorylation sites mapped on the γ-tubulin as in Figure 6A, illustrating the position of the phosphorylation sites in relation to the interface with α-tubulin, shown as in Figure 2B.

Three sites on Spc98p (510,528,531) map near the γTuSC dimer interface, with two of the sites (528 and 531) mapping near *γ*-tubulin. The only other mapped site is Y438, at the C-terminal end of helix 10 (GCP4 numbering), on the face of the γTuSC that would be internal to an assembled *γ*TuRC (Figure 6). Intriguingly, 11 unmodeled sites on Spc98p are all located in the 50 amino-acids preceding our first modeled N-terminal residue with recent work showing the phosphomimetic mutation S152D leading to a cell cycle delay(Fong et al., 2018).

All but one of the mapped sites in Spc97p are located in positions external to an assembled *γ*TuRC (Figure 6). Sites 84 and 88 are in the N-terminus at the Spc110p binding site, suggesting a role in modulating the interaction with Spc110p during γTuSC recruitment to the SPB. Sites 130, 208 and 209 are all located on loops near the bend in the Spc98p structure near the second and third α-helical bundles. Four mapped sites (130, 208, 209, 224,) are located near the second and third helical bundles. S130 is located adjacent to the interface between Spc97p and Spc98p, while S208 and S209, S224, and the unmodeled S219 are located on an insertion between helices 6 and 7 (GCP4 numbering) with S208 and S209 immediately C-terminal to helix 6. Y429 is on a loop buried at the interface between Spc97p and *γ*-tubulin, sterically inaccessible to kinases in our structures. S797 is located near the dimer interface between Spc97p and Spc98p. Recent work has shown that mutations of S84, S88 and S797 have mild effects on spindle assembly, consistent with their playing a role in the regulation of *γ*TuRC assembly and activation(Fong et al., 2018). However, the limited effect suggests that other factors likely also play significant roles.

*γ*-Tubulin is phosphorylated at 14 or 15 sites. A cluster of 6 or 7 sites (S37, S42, T44 and possibly S43 on the H1-S2 loop, and S360, Y362 on the S9-S10 loop) is found near the Spc97,98p:*γ*-tubulin interface and the fourth C-terminal bundle, and near the active (closed) *γ*-tubulin:*γ*-tubulin interface, suggesting a role in modulating *γ*-tubulin interactions with itself or the GCPs during assembly and activation. Consistent with this hypothesis, phosphomimicking mutations of S360 cause a cell cycle delay and defects in spindle morphology(Lin et al., 2011). T130 on the H3-S4 loop may play a similar role, as it also localizes to Spc97,98p:*γ*-tubulin and *γ*-tubulin:*γ*-tubulin interfaces. S373 and the ambiguously assigned S378/S381 sites are located on S10, with S381 being the only site that appears sterically accessible in our structures, on the S10-H11 loop. S381, together with S444 and Y445 are located in a cluster near the C-terminus, potentially modulating the interaction of the C-terminus with the GCPs. *γ*-tubulin Y445 makes a structurally conserved interaction with an arginine on both GCPs (R627 on Spc97p and R633 on Spc98p (Figure 6B)). Phosphomimicking (Y445D) mutations at this site yield a phenotype identical to deletion of the C-terminal conserved DSYL residues, resulting in cell cycle arrest prior to anaphase, as well as effects on microtubule nucleation and spindle organization(Vogel et al., 2001). It is possible that this structurally conserved contact is abrogated by phosphorylation given the highly charged environment of the *γ*-tubulin:GCP interface near *γ*-tubulin’s C-terminus (Figure 3C), modulating the *γ*-tubulin conformation.

Finally S71, T227, Y407 and S415 localize near the *γ*-tubulin:α-tubulin interaction, and are likely to play a role in modulating the affinity of this interaction.

## Discussion

Using a single-particle cryo-EM approach, we have determined structures for monomeric and dimeric *γ*TuSCs at near-atomic resolution. We observe that in contrast to all previous structures which were of free *γ*-tubulin and showed a bent-like conformation, *γ*-tubulin adopts a “straight-like” conformation upon assembly into the *γ*TuSC. In the case of *γ*-tubulin, these conformational changes are allosterically driven by binding to Spc97p or Spc98p. Modeling the γTuSC-bound *γ*-tubulin onto the minus end of an α-tubulin within a yeast microtubule showed that the observed conformational changes would effectively eliminate areas of potential steric clashes, thereby activating the *γ*-tubulin for nucleation. Further, the changes in *γ*-tubulin structure are strikingly similar to the bent-straight transformations that occur in αβ-tubulin upon assembly into the microtubule lattice, suggesting the generality of assembly-driven allosteric activations within the tubulin family.

The near-straight conformation of *γ*-tubulin within monomeric *γ*-TuSC suggests that no significant other conformational changes within the *γ*-tubulin itself would be necessary to optimize its ability to nucleate MTs. Rather, it is the position of the *γ*-tubulins within the *γ*TuRC ring that needs to be allosterically optimized. These conformational changes are likely to be primarily driven by PTMs modulating the interaction of *γ*TuSC with Spc110p, or the allosteric formation of an activated closed *γ*TuRC by affecting the conformations of Spc97p or Spc98p.

Our single particle structures of monomers and dimers all show that Spc97p and Spc98p adopt an open conformation at the intra-TuSC interface. The conformational changes observed in Spc97p and Spc98p upon assembly into dimers suggest that twisting of their α-helical bundles drives changes in the location and orientation of their bound *γ*-tubulins and that these represent a second level of assembly driven-activation. Fitting our structures into the previously determined open and closed assembled *γ*TuRCs shows that Spc98p undergoes the largest conformational changes as it transitions from the open to the closed state during activation, suggesting that either PTMs or the binding of other factors such as Hrr25 (Peng et al., 2015) could be the main allosteric drivers for the formation of a closed state.

Our structures complement existing structural and biochemical data with a high-resolution snapshot of the *γ*TuSC. Together with previous work, these provide a framework for understanding the molecular basis for MT nucleation and regulatory processes likely necessary to ensure that microtubules are only nucleated at the SPB. Our work suggests that functional nucleation is positively controlled in at least three ways: i) the newly discovered changes in *γ*-tubulin upon assembly into *γ*TuSC, ii) assembly of *γ*-TuSCs into and open ring mediated by Spc110p oligomers, and ii) closure of each γTuSC from an open state to a closed state to fully align the *γ*-tubulins to the MT lattice. Further regulation could occur by PTMs at the *γ*:α-tubulin interface that affect *γ*-tubulin’s affinity for αβ-tubulin, but not its conformation. Significant gaps remain in our understanding of the how binding of regulatory proteins and PTMs act to modulate the assembly of the *γ*-TuRC, its interactions with αβ-tubulin, and the formation of its active state.

## Acknowledgements

We gratefully acknowledge many helpful discussions with members of the Agard lab, as well as with our collaborators on the Yeast Centrosome - Structure, Assembly, and Function program project grant in the labs of Trisha Davis, Chip Asbury, Ivan Rayment, Andrej Sali, Mark Winey, and Sue Jaspersen. We would like to thank Michael Braunfeld, Alexander Myasnikov, and David Bulkley for their work running the electron microscopy facility at UCSF, Cameron Kennedy, Matthew Harrington and Joshua Baker-Lepain for their work running the UCSF MSG and wynton clusters, Andrew Lyon and Michelle Moritz for help and training with the purification of *γ*TuSC, and Ray Yu-Ruei Wang for advice with modelling in Rosetta. We acknowledge financial support from the following sources: Howard Hughes Medical Institute (DAA), National Institute of General Medical Sciences (NIGMS) grants: R01 GM031627, R35GM118099 and P01 GM105537.

## Conflicts of Interest

The authors declare that they have no conflicts of interest.

## Materials and Methods

### γTuSC Purification

γTuSC was prepared essentially as described (Vinh et al, 2002, Lyon et al, 2015).

### Grid Preparation

All steps were performed on ice, or in a cold room at 4C. Prior to grid preparation *γ*-TuSC aliquots were centrifuged in a benchtop centrifuge (Eppendorf 5415D) at 16’000 g for 15 minutes, and transferred to a new tube. The sample concentration was assessed on a nanodrop, and diluted to a final concentration of ~1 μM (O.D. at 280nm wavelength of 0.28-0.35) such that the final buffer conditions were 40 mM HEPES pH 7.5, 2 mM MgCl2, 1 mM EGTA, 1 mM GDP, 100 mM KCl and 2.5% v/v glycerol.

Data used for initial model generation and refinement had final buffer conditions of 40 mM HEPES pH 7.5, 1 mM MgCl2, 1 mM EGTA, and 100 mM KCl.

C-flat 1.2-1.3 4C grids were used for sample freezing and glow discharged for ~30s at -20 mA immediately prior to plunge-freezing. Grids were frozen on a Vitrobot Mark II or Mark IV, with the humidity set to 100%, and using Whatman 1 55 mm filter papers.

### Electron Microscopy

Micrographs used in initial model generation were collected using an FEI Tecnai F20 operated at 200 kV at a nominal magnification of 29’000X (40’322 at the detector). The data was collected with a 20 um C2 aperture, and a 100 micron objective aperture with a target underfocus of ~1-2.5 microns. UCSF Image4(Li et al., 2015) was used to operate the microscope. Dose-fractionated micrographs were collected on a Gatan K2 Summit camera in super-resolution mode at a dose rate of ~8.5-9.5 electrons per physical pixel per second for 12 seconds, with the dose fractionated into 40 frames.

Micrographs included in the final model were collected using an FEI Tecnai Polara operated at 300 kV at a nominal magnification of 31’000X (39’891X at the detector). Data was collected with a 30 um C2 aperture and a 100 um objective aperture inserted with a target underfocus of ~1-3 microns. Leginon (Suloway et al., 2005) or SerialEM (Mastronarde, 2005) were used to operate the microscope. Dose-fractionated micrographs were collected on a Gatan K2 Summit camera in superresolution mode at a dose rate of ~6 electrons per physical pixel per second for 20 seconds, with the dose fractionated into 100 frames.

### Image Processing - Initial model generation

Dose-fractionated image stacks were corrected for drift and beam-induced motion as well as binned 2-fold from the super-resolution images using MotionCorr(Li et al., 2013). CTF estimation was performed using CTFFIND4(Rohou and Grigorieff, 2015). Particle coordinates were semi-automatically picked from filtered and binned images using the e2boxer “swarm” tool(Tang et al., 2007). Particles were extracted using Relion(Scheres, 2012) with a box size of 384 physical pixels resampled to 96 pixels for initial processing. A dataset of ~50’000 particles from 217 micrographs was used to generate 300 2D classes using Relion 1.3. 23 classes were selected and used in the generation of a *γ*-TuSC monomer initial model using the e2initialmodel.py function in EMAN2. This model was then used as a reference in Relion 1.3 for 3D classification into 4 classes of a 115’701 particle dataset from 507 micrographs with a 384 pixel box. Particles from the best *γ*-TuSC monomer class were then used for further processing and classification into 4 classes in Frealign(Grigorieff, 2016). The best class, with a resolution of ~9 Angstroms was then used as a 3D reference for processing of the Polara data.

### Image processing - Polara Data

Images were drift-corrected and dose-weighted using MotionCor2(Zheng et al., 2017). CTF estimation was performed using CTFFIND4. Particle coordinates were semi-automatically picked from 6826 filtered and binned images using the e2boxer “swarm” tool, resulting in an initial dataset of ~1.35 million particle coordinates. Particles were extracted from unbinned super-resolution micrographs using Relion’s extraction tool with a box size of 401.088 Å (640 unbinned pixels). Two rounds of 2D classification were performed on images Fourier cropped to 64×64 pixels in Relion 1.4 to eliminate junk particles, resulting in an 883’846 particle dataset. An initial 3D classification into 6 classes was performed in Relion 1.4 with data Fourier cropped to 64×64 pixels, using a centered monomer reconstruction obtained from preliminary TF20 data as described above, resampled to match the Polara data pixel size. The particles and alignment parameters, but not the class assignments were then exported to Frealign, where all further processing was performed. An initial 3D classification into 20 classes to resolve compositional heterogeneity and eliminate residual “junk” particles was performed with images resampled to 64 pixels. γTuSC monomer and dimer classes were separately pooled prior to alignment against a single class. Monomer particles (448’490 particles) - Fourier cropped to 384 pixels in Relion - were aligned against a single reference prior to classification into 3 classes.

Dimer particles (148’622 particles) - initially Fourier cropped to 128 pixels, then finally to 256 pixels - were aligned against a single reference prior to classification into 2 classes.

### Masked reconstructions

Masks were generated from preliminary atomic models of γTuSC, binarized using the e2proc3d.py from the EMAN2 image processing package, and supplied as binary masks to Frealign. A 4 pixel soft cosine edge was applied to the masks in Frealign, with density outside the mask being filtered to 18A resolution with a 3 pixel cosine edge. The full monomer particle stack was used for refinement into a single class.

### Atomic Model Generation - Monomers

To generate an atomic model, the crystal structure of human GCP4 and a previously generated pseudo-atomic model were used as templates. Prior to fitting, the GCP4 structure was threaded with the Spc97p and Spc98p sequence, and the human *γ*-tubulin was threaded with the Tub4p sequence. These initial models were fitted into preliminary structures into segmented density using Rosetta’s relax function. Missing residues were built using RosettaCM density-guided model building(DiMaio et al., 2015), with the human GCP4, *γ*-tubulin threaded models and the pseudo-atomic model being sampled separately. Well scoring structures were then compared to the density, assessing the quality of the fit to determine the register. In cases where the register was poorly fit and the correct register was clear, the register was manually adjusted to fit map details. Certain regions were built using the RosettaES algorithm(Frenz et al., 2017). This procedure was iterated, with occasional manual modification of the structure in Coot.(Emsley et al., 2010)

As a final step, final half-maps were used in the refinement, with the best preliminary models were relaxed and refined through iterative backbone rebuilding (Wang et al., 2016) into one half-map reconstruction, and iteratively refined using Rosetta. Final refinements were performed with GDP density masked out, and GDP was manually docked into its density.

### Model Generation - Dimer

The dimer model was generated by using Rosetta’s relax function to fit two monomer models generated as above into dimer density with GDP masked out.

### Model Generation - Open and Closed Fitting

Monomer models were fitted into segmented open and closed density maps using Rosetta’s relax function. The pixel size for the reconstructions was optimized by fitting models into densities where the pixel size had been changed, followed by visual inspection of the resulting models.

### 2D Classification - Figure S1

Monomer and dimer stacks (384 pixel stacks used in final reconstruction generation) were separately classified using cisTEM (Grant et al., 2018). Classes showing high resolution features were extracted for figure generation using IMOD(Kremer et al., 1996).

### Figure Generation

Structural figures were generated in UCSF Chimera. FSC plots were generated in Excel from Part_FSC estimates in Frealign. Figures panels were compiled into figures in Affinity Designer.

